# Dynamics of host glutathione and glutathione related enzymes in *Macrophomina phaseolina*-*sorghum bicolor* interaction

**DOI:** 10.1101/853986

**Authors:** Ananda Y. Bandara, Dilooshi K. Weerasooriya, Sanzhen Liu, Christopher R. Little

**Affiliations:** Department of Plant Pathology, Kansas State University, Manhattan, KS, 66506, USA.,, and; Department of Agronomy, Kansas State University, Manhattan, KS, 66506, USA

**Keywords:** *Sorghum bicolor*, *Macrophomina phaseolina*, glutathione, glutathione S-transferase, glutathione peroxidase, glutathione reductase

## Abstract

Glutathione and its related enzymes play an integral role in cellular detoxification processes and redox buffering. A genome wide transcriptome profiling was conducted through RNA sequencing to investigate the dynamics of glutathione and related enzymes in *sorghum bicolor* (L.) Moench in response to *Macrophomina phaseolina* (MP) infection. Compared to mock inoculated control treatment, MP significantly upregulated the glutathione synthetase, glutamate cysteine ligase (involved in glutathione biosynthesis), glutathione s-transferase (GST), glutathione peroxidase (GPX), and glutathione reductase (GR) genes in a charcoal rot susceptible sorghum genotype (Tx7000), but not in a resistant genotype (SC599) at 7 days post- inoculation. The net log2 fold up-regulation of the aforesaid genes in MP-inoculated Tx7000 was 1.9, 0.9, 120.0, 9.0, and 4.5, respectively. To confirm the gene expression data, cell extracts were acquired from MP- and mock-inoculated resistant (SC599, SC35) and susceptible (Tx7000, BTx3042) sorghum genotypes and their reduced (GSH), oxidized (GSSG) glutathione, GST, GPX, and GR activities were measured using standard protocols. A significantly reduced GSH/GSSG ratio was observed in Tx7000 and BTx3042 indicating the strong oxidative stress experienced by charcoal rot susceptible genotypes under MP infection. MP significantly increased the GST, GPX, and GR activities of Tx7000 and BTx3042. Although augmented GR activity contributes to cellular GSH restoration, the enhanced GST activity leads to diminishing GSH pools through vacuolar sequestration of GSH-S-conjugates. This eases the oxidative stress confronted by susceptible genotypes under MP infection and in turn contributes to reduced charcoal rot susceptibility. The importance of GSH in controlling the MP infection associated oxidative stress was further supported by the significantly reduced disease severity observed in Tx7000 and BTx3042 upon exogenous GSH application.

## INTRODUCTION

Glutathione, the tripeptide γ-glutamyl-cysteinyl-glycine, plays a key role in detoxification and redox buffering processes in the cell (Noctor and Foyer, 1998). It is the most abundant form of organic sulphur in plants (Dixon et al., 1998). Reduced glutathione (GSH) is the most vital intracellular non-protein thiol compound and plays a major role in the protection of cell and tissue structures from oxidative injury (Foyer and Noctor 2005; Foyer and Noctor 2009; Foyer and Noctor 2011). Among others, such as vitamin C, vitamin E, plant polyphenols, and carotenoids, GSH is a key non-enzymatic antioxidant (Shahidi and Zhong, 2010). These non-enzymatic antioxidants neutralize reactive oxygen species (ROS) through a process known as radical scavenging (Nimse and Pal, 2015). Within cells, free glutathione is mainly present in its reduced form (GSH), which could be rapidly oxidized to glutathione disulfide (GSSG) under oxidative stress. Therefore, the GSH to GSSG ratio is an informative indicator of oxidative stress (Marí et al., 2009). Plants respond to pathogen attacks by varying the levels of GSH. For instance, an increase in GSH content has been reported in leaves attacked by avirulent biotrophic pathogens (Edwards et al., 1991; El-Zahaby et al., 1995; Vanacker et al., 1998) while a decrease has been reported in leaves attacked by some necrotrophic fungi (Gonnen and Schlösser, 1993; Kuzniak and Sklodowska, 1999). GSH is synthesized from amino acids by the sequential action of *g-glutamylcysteine synthetase* (*glutamate cysteine ligase*) and *glutathione synthetase* (Alscher and Hess, 1993). The de-novo synthesis of glutathione from its amino acid constituents is required for the elevation of glutathione as an adaptive response to oxidative stress (Nimse and Pal, 2015).

Glutathione S-transferase (GST) is an important antioxidant enzyme which catalyzes the conjugation of GSH to an electrophilic substrate (Edwards et al., 2000). Many secondary metabolites produced by plants are phytotoxic even to the cells that produce them, and therefore the appropriate cellular localization (usually the vacuole) is important (Matern et al., 1986; Sandermann, 1992; Sandermann, 1994) and the GSH/GST system plays a key role in phytotoxin compartmentalization. For example, anthocyanin pigments require GSH conjugation by GST for transport into the vacuole as inappropriate cytoplasmic retention of anthocyanins leads to cytotoxicity (Marrs et al., 1995). Moreover, the endogenous products of oxidative damage initiated by reactive oxygen species such as lipid peroxides (e.d. 4-hydroxyalkenals) and oxidative DNA degradation products (e.g. base propanols) are cytotoxic. Plant and animal GSTs play a key role in conjugating GSH with such endogenously produced electrophiles, which results in their detoxification (Bartling et al., 1993; Berhane et al., 1994; Danielsonn et al., 1987). Previous findings showed that the transcription of plant GST genes is regulated by various abiotic (Edwards et al., 2000; Seppanen et al., 2000; Moons, 2003; Kiyosue et al., 1993; Bianchi et al., 2002) and biotic stresses such as pathogen attack (Mauch and Dudler, 1993; Liao et al, 2014).

Glutathione peroxidase is another major ROS scavenging enzyme in plants (Mittler et al., 2004). Expression of glutathione peroxidase has been found to be highly up-regulated in response to pathogen infection (Agrawal et al., 2002; Levine et al., 1994). Using reduced glutathione (GSH) as an electron donor, it catalyzes the reduction of H_2_O_2_ or organic hydroperoxides to water or corresponding alcohols while GSH is oxidized into glutathione disulfide (GSSG) (Margis et al., 2008). GSSG is reduced back to GSH by glutathione reductase in an NADPH-dependent manner (Meloni et al., 2003; Huber et al., 2008). Glutathione reductase secreted by *Magnaporthe oryzae* has been shown to be required for neutralizing plant generated ROS during the rice blast disease (Fernandez and Wilson, 2014).

*Macrophomina phaseolina* is a soilborne, necrotrophic fungal pathogen that causes diseases in over 500 different plant species (Islam *et al*., 2012). It causes charcoal rot disease in many economically important crops such as sorghum, soybean, maize, alfalfa and jute (Islam *et al*., 2012). Charcoal rot is a high priority fungal disease in sorghum [*Sorghum bicolor* (L.) Moench], causing tremendous crop losses where ever sorghum is grown (Tarr, 1962, Tesso *et al*., 2012). Recent reports revealed the *M. phaseolina*’s ability to negatively affect the physicochemical properties of sorghum grain (Bandara et al., 2017 a), sorghum yield components (Bandara et al., 2017 b), leaf greenness (Bandara et al., 2016) and sweet sorghum biofuel traits such and juice yield and biomass (Bandara et al., 2017 c). Our recent genes expression and functional investigations provided exciting insights into induced charcoal rot disease susceptibility in grain sorghum through up-regulated host oxidative stress after *M. phaseolina* infection (findings have been submitted for publication). In this context, the dynamics of host glutathione and its related enzymes are worthy of study to gain insights into potential role of glutathione system in coping with enhanced host oxidative stress under *M. phaseolina* infection. Therefore, the objectives of the current study were (i) to discover differentially expressed glutathione related genes between known charcoal rot resistant and susceptible sorghum genotypes in response to *M. phaseolina* inoculation and (ii) to measure the glutathione (reduced and oxidized) concentration and activity of glutathione related enzymes (glutathione-s transferase, glutathione peroxidase, glutathione reductase) in *M. phaseolina*-infected stalk tissues. Therefore, our overarching goal here is to uncover the potential links between glutathione (and related enzymes) and charcoal rot disease reaction at transcriptional and biochemical levels.

## RESULTS

### Differential expression of genes related to metabolism of host glutathione and its related enzymes in response to *M. phaseolina* infection

Supplementary table 1 provides a summary of the differentially expressed genes between two sorghum genotypes upon *M. phaseolina* inoculation that are related to metabolism of host glutathione and its related enzymes. Differential gene expression analysis revealed eleven and 52 glutathione related genes that are differentially expressed between SC599 and Tx7000 in response to pathogen inoculation at 2 and 7 DPI, respectively (Figure 1). None of the glutathione related genes were differentially expressed at 30 DPI. Out of eleven differentially expressed genes at 2 DPI, eight encoded GST, two encoded Glutaredoxin while the remaining gene encoded GPx. There was a net GST transcripts upregulation (log2 fold = 4.16) in Tx7000 after pathogen inoculation. Both Glutaredoxin genes were significantly down-regulated in SC599 (net log2 fold = −2.18) while one of them was significantly up-regulated in Tx7000 (net log2 fold = 1.27). The GPx gene was significantly up-regulated in SC599 while that of Tx7000 did not change. Out of 52 glutathione-related differentially expressed genes at 7 DPI, 41 encoded GST, six GPx, one each for glutathione synthetase and glutamate cysteine ligase, and two each for GR and glutaredoxin. Out of the 41 GST genes, 32 were significantly up-regulated in pathogen-inoculated Tx7000 while six were significantly down-regulated. The majority (except three genes) of these genes in SC599 were not significantly differentially expressed. The net log2 fold up-regulation of GST genes in pathogen-inoculated Tx7000 was 120 while the net log2 fold down-regulation of GST genes in pathogen-inoculated SC599 was 11.6. Out of five GPx genes, four were significantly up-regulated in pathogen-inoculated Tx7000 while one was significantly down-regulated. The net log_2_ fold up-regulation was 9. None of these genes were significantly differentially expressed in SC599. Both glutathione synthetase and glutamate cysteine ligase genes were significantly up-regulated in pathogen-inoculated Tx7000 while those of SC599 were non-significantly down-regulated. The two GR genes were significantly up-regulated in pathogen-inoculated Tx7000 (net log2 fold = 4.5) while the two glutaredoxin genes were significantly down-regulated (net log2 fold = 4.6). None of these four genes were significantly differentially expressed in SC599 after pathogen inoculation.

**Figure 1.**
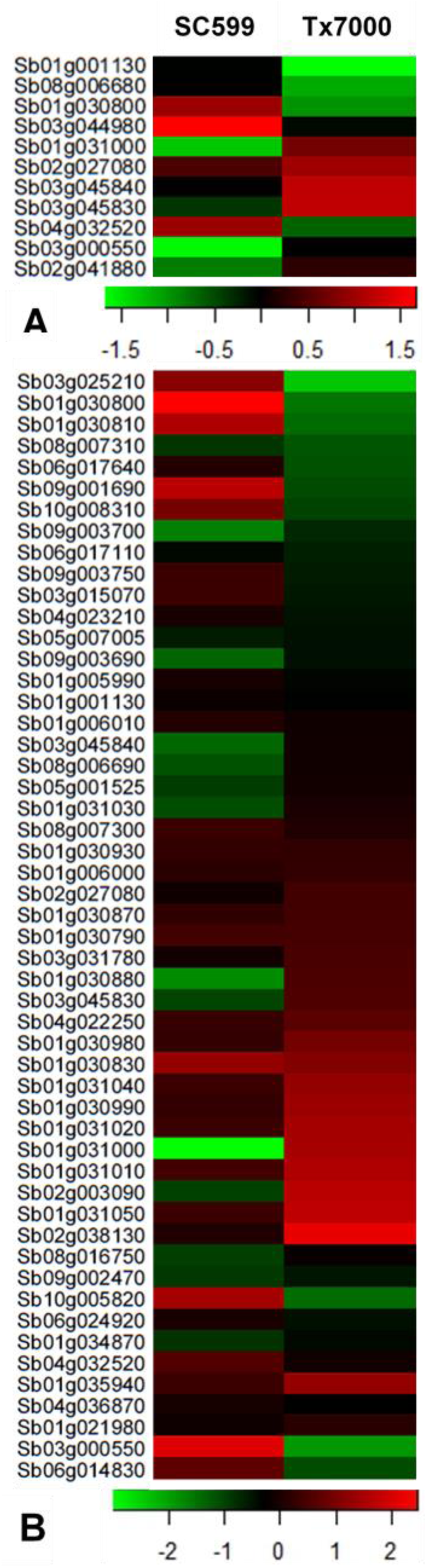
Heat map depicting differentially expressed glutathione-related genes between charcoal rot resistant (SC599) and susceptible (Tx7000) sorghum genotypes in response to *Macrophomina phaseolina* inoculation at 2 (A) and 7 (B) days post-inoculation. Red, green, and black colors respectively represent up-regulated, down-regulated, and non-differentially expressed genes after *M. phaseolina* inoculation in comparison to mock-inoculated control treatment with sterile phosphate buffered saline.

### Analysis of variance (ANOVA) for functional assays and disease severity experiment

Table 1 provides the F-statistic *P*-values from the analysis of variance (ANOVA) for the functional assays conducted in the current study. Although the treatment (*M. phaseolina*- and mock-inoculated control) had a significant main effect on total, oxidized, and reduced glutathione concentration and GPx activity at 4 DPI, treatment effect was genotype-specific for said response variables at 7 and 10 DPI. Treatment did not have a significant main or simple effect on the reduced to oxidized glutathione ratio and GR activity at 4 DPI. However, the genotype by treatment interaction was significant on reduced to oxidized glutathione ratio and GR activity at 7 and 10 DPI. The genotype-by-treatment interaction was significant on GST activity at all post inoculation stages. The genotype-by-treatment interaction was also significant for lesion length (*P* = 0.0089).

**Table 1.**
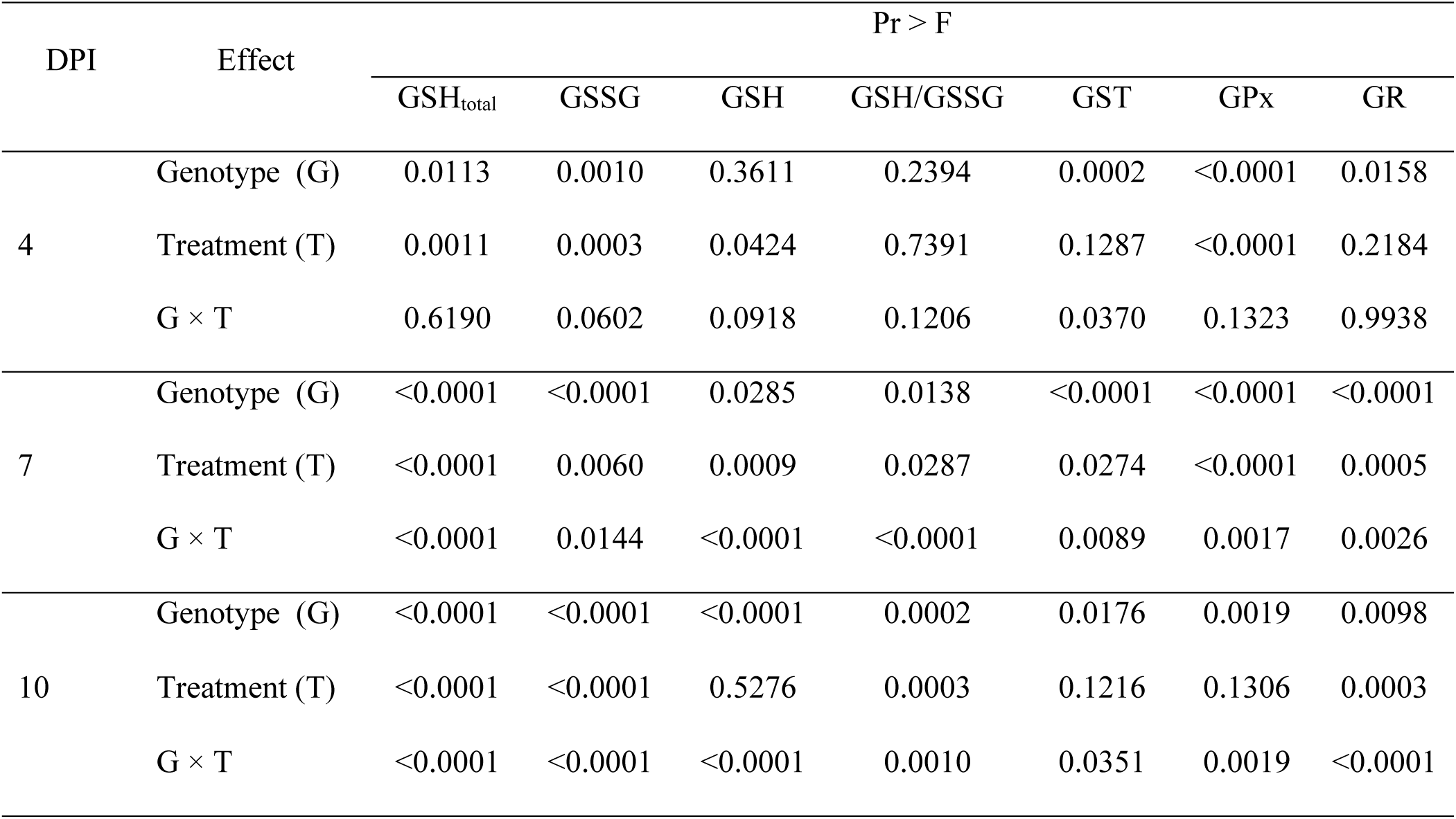
*P*-values of F-statistic from analysis of variance (ANOVA) for functional assays including total glutathione (GSH_total_), oxidized glutathione (GSSG), reduced glutathione (GSH), reduced to oxidized glutathione ratio (GSH/GSSG), glutathione-s-transferase activity (GST), glutathione peroxidase activity (GPx), and glutathione reductase activity (GR) measured with four sorghum genotypes (Tx7000, BTx3042, SC599, SC35) after inoculation with *M. phaseolina* at 3 post inoculation stages (4, 7, and 10 days post-inoculation, DPI) (α = 0.05).

| DPI | Effect | Pr > F |  |  |  |  |  |  |
| --- | --- | --- | --- | --- | --- | --- | --- | --- |
|  |  | GSH<sub>total</sub> | GSSG | GSH | GSH/GSSG | GST | GPx | GR |
| 4 | Genotype (G) | 0.0113 | 0.0010 | 0.3611 | 0.2394 | 0.0002 | <0.0001 | 0.0158 |
|  | Treatment (T) | 0.0011 | 0.0003 | 0.0424 | 0.7391 | 0.1287 | <0.0001 | 0.2184 |
|  | G × T | 0.6190 | 0.0602 | 0.0918 | 0.1206 | 0.0370 | 0.1323 | 0.9938 |
| 7 | Genotype (G) | <0.0001 | <0.0001 | 0.0285 | 0.0138 | <0.0001 | <0.0001 | <0.0001 |
|  | Treatment (T) | <0.0001 | 0.0060 | 0.0009 | 0.0287 | 0.0274 | <0.0001 | 0.0005 |
|  | G × T | <0.0001 | 0.0144 | <0.0001 | <0.0001 | 0.0089 | 0.0017 | 0.0026 |
| 10 | Genotype (G) | <0.0001 | <0.0001 | <0.0001 | 0.0002 | 0.0176 | 0.0019 | 0.0098 |
|  | Treatment (T) | <0.0001 | <0.0001 | 0.5276 | 0.0003 | 0.1216 | 0.1306 | 0.0003 |
|  | G × T | <0.0001 | <0.0001 | <0.0001 | 0.0010 | 0.0351 | 0.0019 | <0.0001 |

### Sorghum glutathione dynamics after *M. phaseolina* inoculation

Compared to control, *M. phaseolina* significantly increased the total (49%, *P* = 0.0011), oxidized (50%, *P* = 0.0002), and reduced (48%, *P* = 0.0424) glutathione concentrations across genotypes at 4 DPI (Figure 2 A, C, E). Although pathogen inoculation significantly reduced the total, oxidized, and reduced glutathione concentration of Tx7000 (40%, *P* < 0.0001; 27.7%, *P* < 0.0001; 58.1%, *P* < 0.0001, respectively) and BTx3042 (43%, *P* < 0.0001; 12.7%, *P* = 0.0471; 81.6%, *P* < 0.0001, respectively) at 7 DPI, inoculation did not significantly affect those in two resistant genotypes, SC599 and SC35 (Figure 2 B, D, F). Interestingly, compared to control, pathogen inoculation significantly increased the total and oxidized glutathione concentration of Tx7000 (161%, *P* < 0.0001; 234%, *P* < 0.0001, respectively) and BTx3042 (192%, *P* < 0.0001; 294%, *P* < 0.0001, respectively) at 10 DPI, although inoculation did not significantly affect the total and oxidized glutathione concentration in SC599 and SC35 (Figure 2 B). Pathogen inoculation significantly decreased reduced glutathione concentration of Tx7000 (36.4%, *P* < 0.0001) while significantly increasing it in BTx3042 (44.6%, *P* < 0.0001) at 10 DPI (Figure 2 F). Inoculation did not significantly affect the reduced glutathione concentration of two resistant genotypes (Figure 2 F). Although pathogen inoculation significantly decreased the reduced to oxidized glutathione ratio of Tx7000 (7 DPI: 41.4%, *P* < 0.0001; 10 DPI: 57.9%, *P* < 0.0001) and BTx3042 (7 DPI: 79.2%, *P* < 0.0001; 10 DPI: 64.6%, *P* < 0.0001) at 7 and 10 DPI, inoculation did not significantly affect the reduced to oxidized glutathione ratio in SC599 and SC35 (Figure 3).

**Figure 2.**
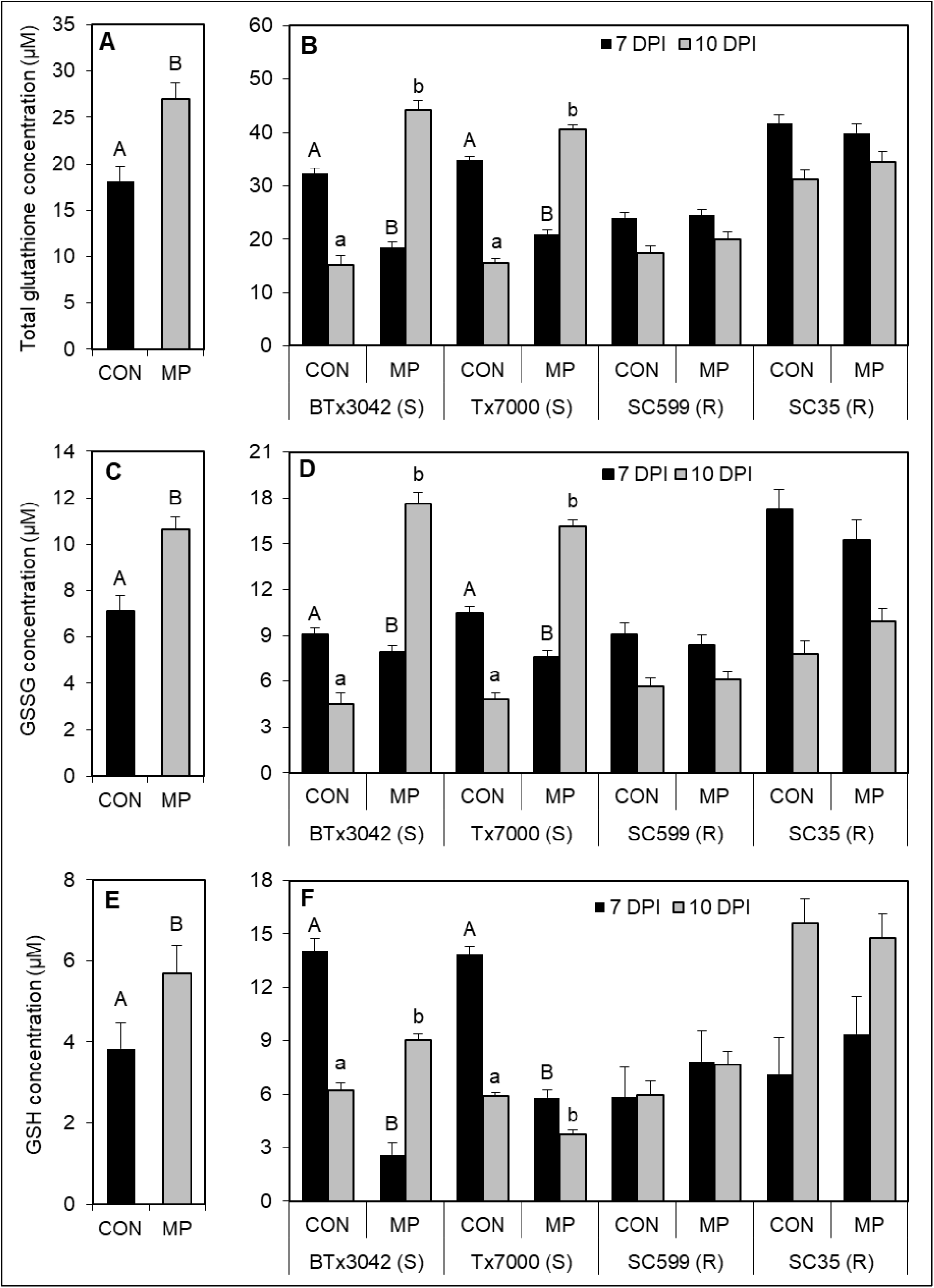
Comparison of the mean total glutathione content between two treatments (A) across four genotypes at 4 days post-inoculation (DPI), (B) among four genotypes at 7 and 10 DPI; oxidized glutathione (GSSG) content between two treatments (C) across four genotypes at 4 DPI, (D) among four genotypes at 7 and 10 DPI; and the reduced glutathione (GHS) content between two treatments (E) across four genotypes at 4 DPI, (F) among four genotypes at 7 and 10 DPI. In panels A, C, and E, treatment means followed by different letters are significantly different. In panels B, D, and F, treatment means followed by different letters within each genotype at a given DPI are significantly different. Treatment means without letter designations within each genotype at a given DPI are not significantly different (α = 0.05). Error bars represent standard errors. CON = phosphate-buffered saline mock-inoculated control, MP = *Macrophomina phaseolina*.

**Figure 3.**
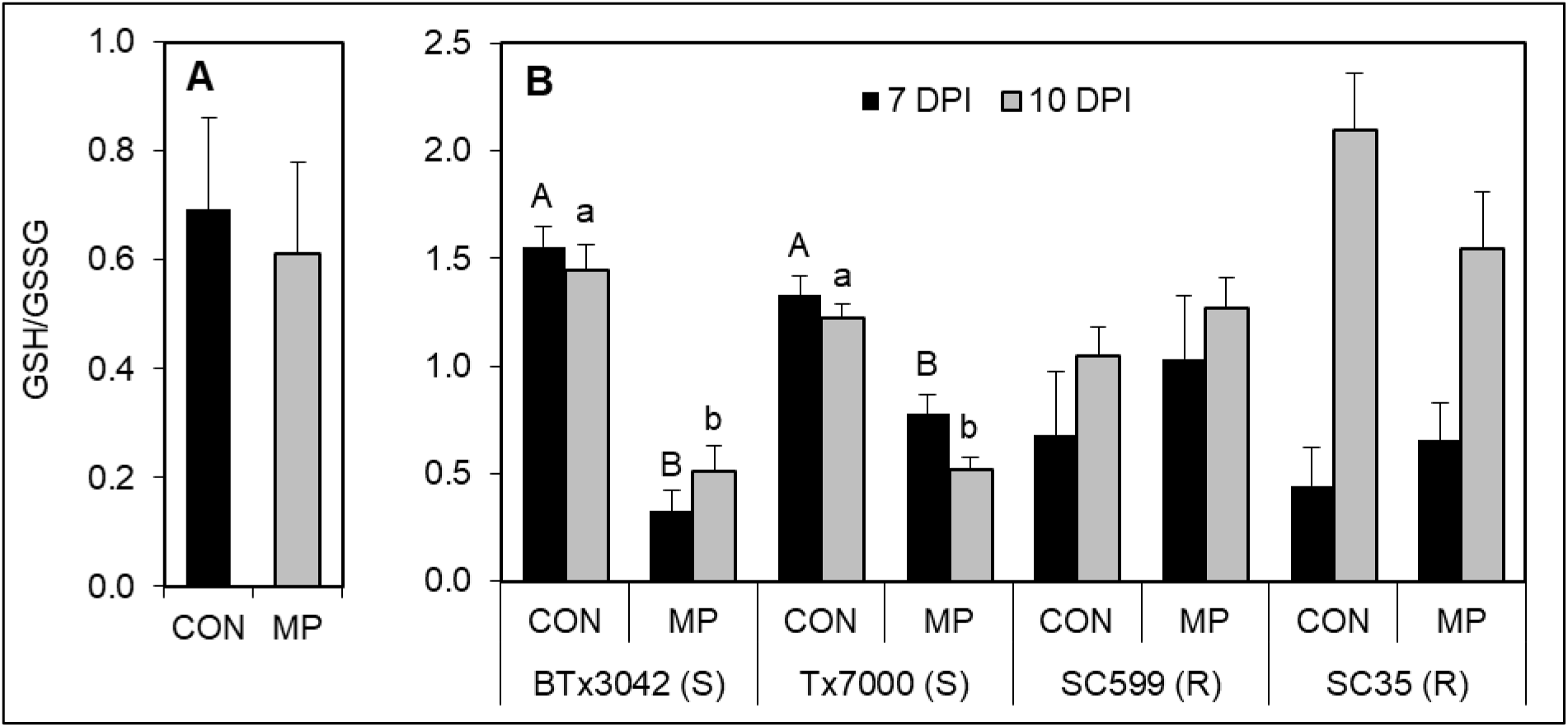
Comparison of the mean GSH/GSSG ratio between two treatments (A) across four genotypes at 4 days post-inoculation (DPI), and (B) among four genotypes at 7 and 10 DPI. Treatment means without letter designations are not significantly different. In panel B, treatment means followed by different letters within each genotype at a given DPI are significantly different while the treatment means without letter designations within each genotype at a given DPI are not significantly different (α = 0.05). Error bars represent standard errors. CON = phosphate-buffered saline mock-inoculated control, MP = *Macrophomina phaseolina*.

### Dynamics of sorghum GST, GPx, and GR enzymes after *M. phaseolina* inoculation

Compared to control, at all post inoculation stages, *M. phaseolina* significantly increased the GST specific-activity (μmol/mL/min) of Tx7000 (4 DPI: 376.8%, *P* = 0.0002; 7 DPI: 233.8%, *P* = 0.0008; 10 DPI: 223.3%, *P* = 0.0354) and BTx3042 (4 DPI: 55.3%, *P* = 0.0325; 7 DPI: 111.5%, *P* = 0.0469; 10 DPI: 164.2%, *P* = 0.0043) (Figure 4 A). However, pathogen inoculation did not significantly affect the GST-specific activity of two resistant genotypes at any post-inoculation stage. Compared to control, pathogen inoculation significantly increased GPx activity (U/L) across genotypes (66.2%, *P* < 0.0001) at 4 DPI (Figure 4 B). Although *M. phaseolina* significantly increased the GPx activity of Tx7000 (7 DPI: 42.5%, *P* < 0.0001; 10 DPI: 35.5%, *P* = 0.011) and BTx3042 (7 DPI: 65.3%, *P* < 0.0001; 10 DPI: 30.8%, *P* = 0.0106) at 7 and 10 DPI, pathogen inoculation did not significantly affect the GPx activity in two resistant genotypes (Figure 4 C). *M. phaseolina* inoculation did not significantly affect the GR activity of tested genotypes at 4 DPI (Figure 4 D). At 7 DPI, pathogen inoculation significantly increased the GR activity of Tx7000 (74.5%, *P* = 0.0363) and BTx3042 (43.2%, *P* < 0.0001). Interestingly, pathogen inoculation significantly reduced the GR activity of Tx7000 (45.4%, *P* < 0.0001) and BTx3042 (29.1%, *P* < 0.0001) at 10 DPI (Figure 4 E). Pathogen did not significantly affect the GR activity of two resistant genotypes at 7 and 10 DPI.

**Figure 4.**
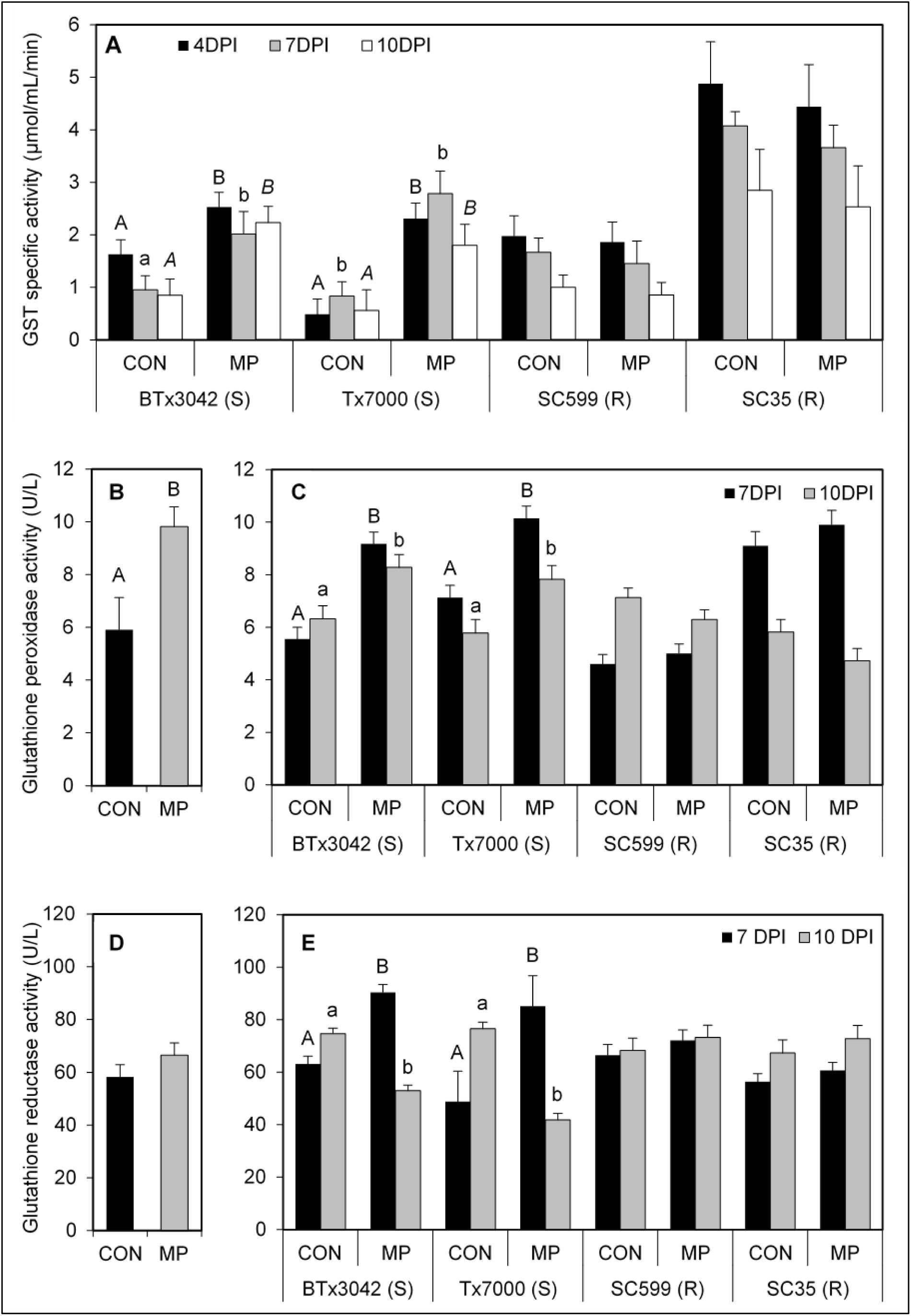
Comparison of the mean (A) glutathione-s-transferase specific activity between two treatments among four genotypes at 4, 7, and 10 days post-inoculation (DPI); glutathione peroxidase activity between two treatments (B) across four genotypes at 4 DPI, (C) among four genotypes at 7 and 10 DPI; and the glutathione reductase activity between two treatments (D) across four genotypes at 4 DPI, (E) among four genotypes at 7 and 10 DPI. In panels B and D, treatment means followed by different letters are significantly different. In panels A, C, and E, treatment means followed by different letters within each genotype at a given DPI are significantly different. Treatment means without letter designations within each genotype at a given DPI are not significantly different (α = 0.05). Error bars represent standard errors. CON = phosphate-buffered saline mock-inoculated control, MP = *Macrophomina phaseolina*.

### Exogenous GSH application reduce charcoal rot disease severity

Compared to the *M. phaseolina* treatment, the *M. phaseolina* + glutathione treatment significantly reduced the lesion length of both charcoal-rot-susceptible genotypes (*P* < 0.016) while glutathione application did not significantly affect the lesion length of the two resistant genotypes (Figure 5).

**Figure 5.**
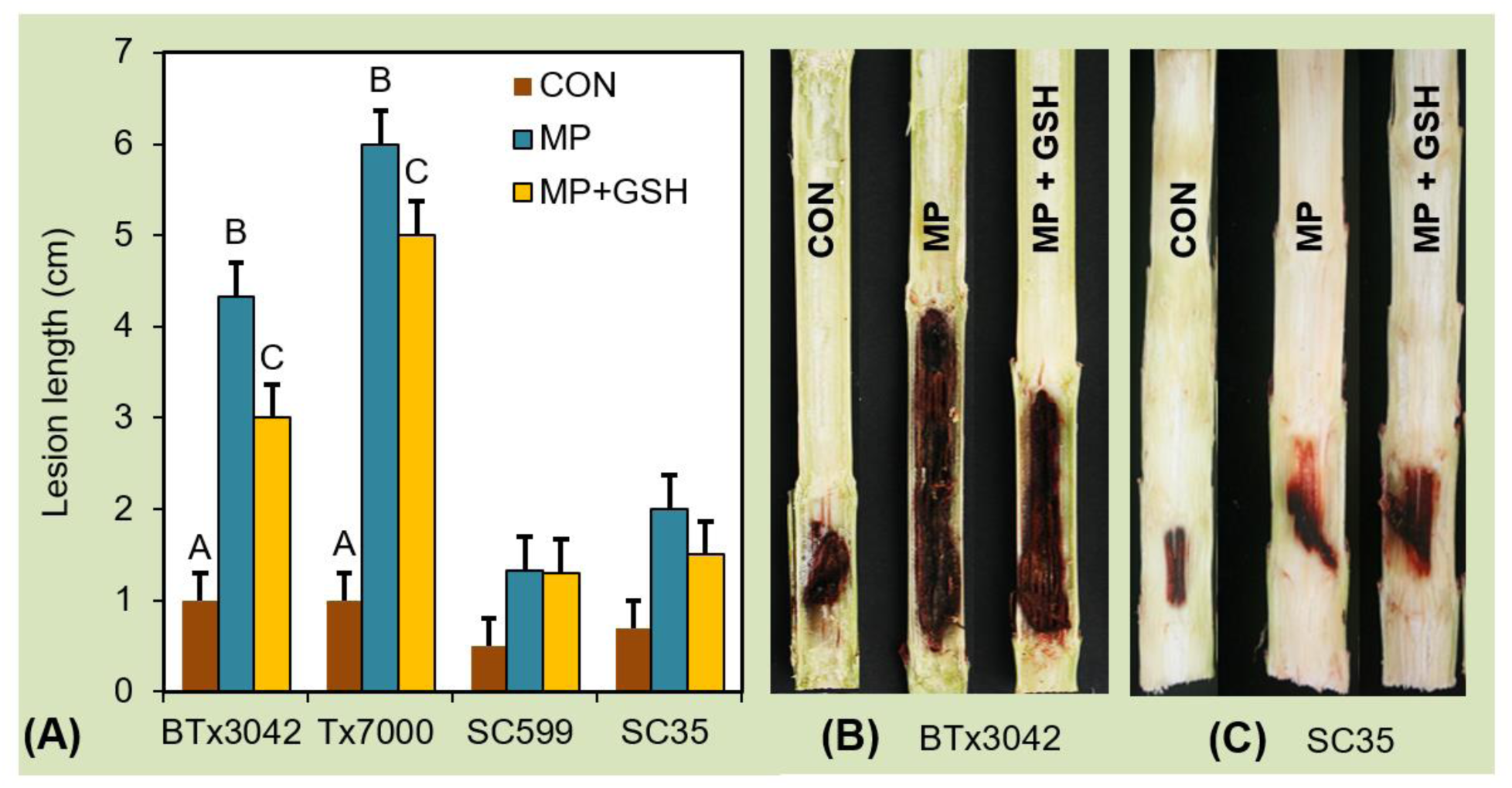
Comparison of mean lesion length between three treatments (CON, MP, MP + GSH) among tested sorghum genotypes at 35 d after inoculation (A). Treatment means followed by different letters within each genotype are significantly different based on the adjusted *P*-value for multiple comparisons using Tukey-Kramer’s test at comparisonwise error rate (α_CER_) = 0.016. The means without letter designations within each genotype are not significantly different. Error bars represent standard errors. CON = phosphate-buffered saline mock-inoculated control, MP = *Macrophomina phaseolina*, GSH = reduced glutathione. Figures B and C show the visual differences between lesion lengths observed in longitudinally split stalks of the charcoal rot susceptible (BTx3042) and resistant (SC35) sorghum genotypes after receiving different inoculation treatments.

## DISCUSSION

Glutathione plays a crucial role in protecting plants from numerous environmental stresses, including oxidative stress due to the generation of active oxygen species, xenobiotics, and some heavy metals (Xiang and Oliver, 1998). In the current study, most of the glutathione-related gene differential expression occurred at 7 DPI revealing the importance of pathogen mediated expression differences of said genes at 7 DPI. Gene expression data at 7 DPI revealed enhanced glutathione biosynthetic capacity; enhanced GST, GPx, and GR activity; and impeded glutaredoxin activity in the charcoal-rot-susceptible sorghum genotype, Tx7000, after *M. phaseolina* inoculation. An increase in the expression of GST and GPx has been identified in soybean cells adjacent to those undergoing the hypersensitive cell death induced by an avirulent phytopathogen (Levine et al., 1994).

Our previous transcriptional and functional investigations revealed the enhanced reactive oxygen/nitrogen species biosynthesis in *M. phaseolina*-inoculated charcoal rot susceptible sorghum genotypes such as Tx7000 and BTx3042. Glutathione is involved in quenching reactive oxygen (Foyer et al., 1994) and nitrogen (Airak et al., 2011) species. Therefore, enhanced glutathione expression helps relaxing the strong oxidative stress encountered by Tx7000 due to pathogen inoculation. Figure 6 depicts the proposed glutathione-mediated antioxidative machinery of charcoal-rot-susceptible sorghum genotype, Tx7000 after *M. phaseolina* infection.

**Figure 6.**
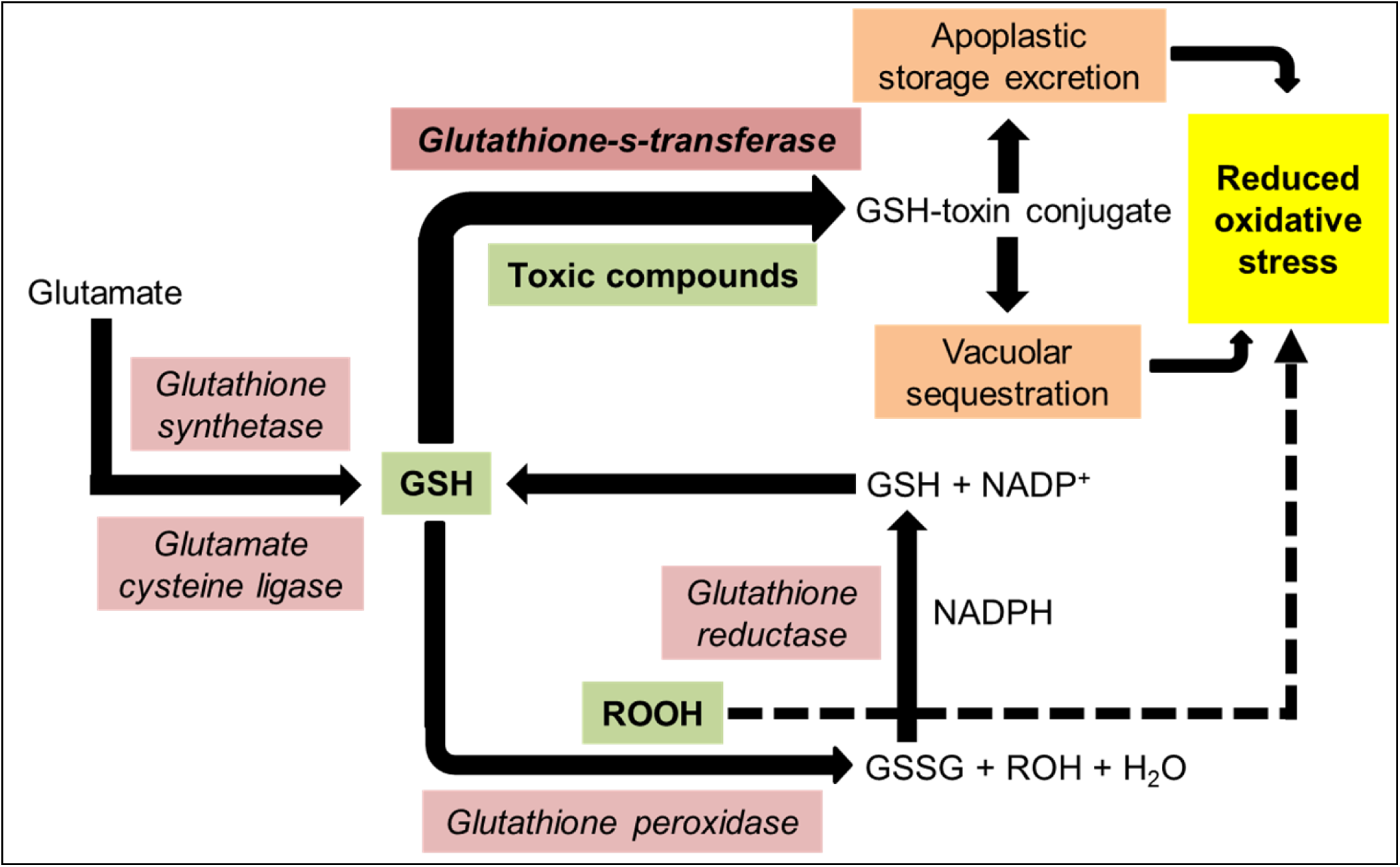
Proposed cellular antioxidative mechanism of charcoal-rot-susceptible sorghum genotype, Tx7000 after *M. phaseolina* infection. GSH = reduced glutathione, GSSG = oxidized glutathione, ROOH = hydroperoxide, ROH = alcohol. *Pink box* = up-regulation/increased activity, *green box* = reduced quantity. Transcriptional and functional data suggested a general enhancement of Tx7000 antioxidative machinery to impede the strong oxidative stress after *M. phaseolina* infection.

To understand the translational aspects of gene expression data in detail, we conducted all functional experiments at three post-inoculation stages (4, 7, and 10 DPI). Functional assays revealed significantly decreased total glutathione, GSSG, and GSH concentrations of two susceptible genotypes at 7 DPI upon pathogen inoculation. Reduced GSH content has previously been observed in tomato leaves infected with the necrotrophic fungus *Botrytis cinerea* (Kuzniak and Sklodowska, 1999) and in *Avena sativa* leaves inoculated with *Drechslera avenae* and *D. siccans* (Gonnen and Schlösser, 1993). Decrease in GSH impedes the host antioxidant capacity and can in turn promote host cell death that facilitates the spread of necrotrophic phytopathogens. GSH is synthesized (*de novo*) from amino acids by the sequential action of *g-glutamylcysteine synthetase* (*glutamate cysteine ligase*) and *glutathione synthetase* (Alscher and Hess, 1993) and the *de-novo* GSH synthesis is required for the elevation of GSH levels as an adaptive response to oxidative stress (Nimse and Pal, 2015). The transcriptional data of the current study suggested the enhanced *de novo* GSH biosynthetic capacity in Tx7000 due to up-regulation of *glutathione synthetase* and *glutamate cysteine ligase*. However, confirming the up-regulated GST and GPx gene expression in pathogen-inoculated Tx7000 at 7 DPI, the functional assays provided evidence for enhanced GST-specific activity and GPx activity in both susceptible genotypes (Tx7000, BTx3042), leading to a net decline in GSH. Despite the enhanced GPx activity, GSSG concentration of pathogen-inoculated susceptible genotypes at 7 DPI remained significantly lower mainly due to enhanced GR activity, which rapidly converts GSSG in to GSH. Therefore, the primary cause behind the decreased amounts of reduced GSH in the pathogen-inoculated susceptible genotypes appeared to be the enhanced GST-specific activity.

In plants, glutathione S-conjugates are either sequestered in the vacuole (Coleman et al., 1997; Wolf et al., 1996) or transferred to the apoplast, a process termed “storage excretion” (Sandermann, 1992; Martinoia et al., 1993; Sandermann, 1994). Therefore, GST activity results in irreversible GSH depletion leading to decreased levels of GSH if not to *de novo* GSH biosynthesis. Moreover, jasmonic acid is a potent expression stimulator for genes involved in GSH biosynthesis and recycling, which could possibly lead to boosted GSH levels (Xiang and Oliver, 1998). Our RNA-Seq data (not present here) showed that some key genes involved in jasmonic acid biosynthetic pathway were strongly down-regulated in *M. phaseolina* inoculated Tx7000 at 7 days post inoculation. Therefore, it seems plausible that down-regulated jasmonic acid biosynthesis in charcoal-rot-susceptible sorghum genotypes after *M. phaseolina* inoculation contributes to decreased GSH recycling, which may limit the availability of GSH.

Interestingly, the pathogen-inoculated susceptible genotypes had significantly increased total glutathione and GSSG concentrations at 10 DPI. Enhanced GPx activity along with reduced GR activity contributes to increased GSSG concentration of pathogen-inoculated susceptible genotypes. Moreover, some GSTs can also function as GPx (Bartling et al., 1993; Cummins et al., 1999) which contributes to increased GSSG concentration. As the GST-specific activity of susceptible genotypes were significantly higher after pathogen inoculation, the observed increase in total glutathione of these genotypes becomes possible only when there is strongly enhanced *de-novo* glutathione biosynthesis. This could contribute to the significantly higher GSH concentration of BTx3042. However, the significantly decreased GSH concentration observed in pathogen-inoculated Tx7000 at 10 DPI is possibly due to its greater rate of GSH utilization (demonstrated by increased GST and GPx activities) than the *de novo* glutathione synthesis and recycling.

It has been suggested that the GSH/GSSG ratio is indicative of the cellular oxidative status and redox balance (Droge, 2002; Foyer and Noctor, 2003). Under strong oxidative stress, GSH is rapidly converted into GSSG, which results in a lower GSH/GSSG ratio. The lower GSH/GSSG ratios observed in susceptible genotypes after *M. phaseolina* inoculation at 7 and 10 DPI further confirmed the strong oxidative stress experienced by these genotypes under *M. phaseolina* inoculation.

In maize, inappropriate accumulation of anthocyanins in cytoplasm causes localized necrosis, poor vigor, or even death of plants. Certain GSTs like BZ-2 has been identified to catalyze the formation of anthocyanin-GSH conjugates, which allows transport into vacuoles thus reducing the cytotoxic effects of higher anthocyanin concentrations (Marrs et al., 1995). The charcoal-rot-susceptible sorghum genotypes tested in this study accumulated comparatively greater amounts of anthocyanins than resistant genotypes, which is manifested as longer pigmented lesions within split stems. In fact, the length of this lesion is used as a measure of charcoal rot resistance. Moreover, ROS are claimed to play a critical role as signaling molecules for anthocyanin production (Hatier and Gould, 2008). Our functional investigations (submitted for publication in another journal) revealed the enhanced ROS biosynthesis in *M. phaseolina* inoculated charcoal rot susceptible genotypes, Tx7000 and BTx3042. Therefore, the overaccumulation of anthocyanins in these genotypes after *M. phaseolina* infection is plausible where their GSH/GST system might play a pivotal role in decreasing the cytotoxic effects of anthocyanin overaccumulation. This, in turn could reduce susceptibility to *M. phaseolina*. Therefore, among many possible substrates for GSH/GST system, anthocyanin could to be a major candidate in compatible charcoal rot reactions.

Elevated GSH biosynthetic capacity has been shown to ironically cause increased oxidative stress in transgenic tobacco plants (Creissen et al., 1999). If this is the case with sorghum after *M. phaseolina* infection, enhanced GSH could escalate charcoal rot susceptibility. Necrotrophic pathogens such as *M. phaseolina* benefited from oxidative stress-mediated host cell death. However, the reduced disease severity observed in two susceptible genotypes after exogenous GSH application shows that GSH does not enhance disease susceptibility, but reduces disease severity. In fact, exogenous GSH can mimic fungal elicitors in activating the expression of defence-related genes (Dron et al. 1988) including PATHOGENESIS-RELATED PROTEIN 1 (Gomez et al. 2004), which contributes to reduced disease susceptibility.

## CONCLUSIONS

Owing to its broad host range, wide geographic distribution, and ability to cause a variety of diseases with higher economic significance, *M. phaseolina* is known to be a globally important necrotrophic fungus. However, compared to other necrotrophic pathosystems and virulence mechanisms, less is known about the *M. phaseolina*’s virulence mechanisms. Classically, necrotrophs are thought to kill the host using various phytotoxins, cell wall degrading enzymes, and reactive oxygen species that are secreted into the host tissues. Our recent findings showed that *M. phaseolina*’s is able to manipulate sorghum metabolic pathways that lead to enhanced oxidative stress and thereby induce charcoal rot disease susceptibility in certain sorghum genotypes. Glutathione and its related enzymes such as GST, GR, and GPx are integral components of the antioxidant system of many life forms including plants and play a pivotal role in maintaining the cellular redox balance. As our current transcriptional and functional investigation suggested, the dynamics of glutathione and related enzymes such as GST, GPx, and GR in charcoal rot susceptible sorghum genotypes (Tx7000 and BTx3042) should be viewed as mechanisms leading to reduced oxidative stress thus charcoal rot susceptibility under *M. phaseolina* infection rather than those results in enhanced disease resistance.

## EXPERIMENTAL PROCEDURES

### A) RNA sequencing experiment

**i) Plant materials, establishment and maintenance, inoculation, and stalk tissue collection**

The treatment structure of the experiment was 2×2×3 factorial (factors respectively represent genotype, inoculation treatment, and tissue collecting stage) with three biological replicates per factor combination. The design structure was completely randomized (CRD). One charcoal rot resistant (SC599) and one susceptible (Tx7000) sorghum genotypes were used. Seeds were treated with the fungicide Captan (N-trychloromethyl thio-4-cyclohexane-1,2 dicarboxamide). Three seeds were planted per pot (19 L Poly Tainer) filled with Metro-Mix 360 growing medium (Sun Gro Bellevue, WA, U.S.A). At three weeks after emergence, each pot was thinned to one seedling per pot. Pots were kept in a greenhouse at 25-32°C with a 16-h light/8-h dark photoperiod. Seedlings/plants were maintained according to the procedures explicated by Bandara et al. (2015). A well characterized, highly virulent *M. phaseolina* isolate (acquired from the row crops pathology lab, Department of Plant Pathology, Kansas State University) was used for inoculation. Mycelial fragments were used as the inoculum. Inoculum preparation and concentration adjustment were performed according to the protocols described by Bandara et al. (2015). At 14 days after anthesis, plants were inoculated by injecting 0.1 mL of inoculum into the basal internode of the stalk using a 1 mL, 26 gauge, 1.5 inch needle, sterile surgical syringe. For mock-inoculation (control treatment), phosphate-buffered saline (pH 7.2) was used. At 2, 7, and 30 days post inoculation (DPI), approximately 8-10 cm long stalk piece containing the inoculation point were collected from each inoculated and mock-inoculated plants. Stalk pieces were immediately frozen in liquid nitrogen and subsequently stored at −80°C until RNA is extracted.

**ii) Total RNA extraction, cDNA library preparation, and Illumina sequencing**

Approximately 1 g of stalk tissues (1 cm above the symptomatic region) was ground in liquid nitrogen using a mortar and pestle and resulted tissue powder was used for RNA extraction. Following the manufacturer’s instructions, total RNA was extracted using Triazole reagent (Thermo Scientific, USA). Upon treatment with Amplification Grade DNAse I (Invitrogen Corporation, USA), the quality and quantity of RNA extracts were determined using Nanodrop 2000 instrument (Thermo Scientific, USA). The required RNA concentration (100-200ng/μl) was obtained by diluting the samples with RNase free water. RNA integrity and quantity were analyzed using Agilent 2100 Bioanalyzer (Agilent Technologies Genomics, USA). Following the manufacturer’s protocol (Illumina Inc., USA), 36 cDNA libraries were prepared using the Illumina TruSeq™ RNA sample preparation kit. RNA from biological replicates was not pooled prior to library preparation. Using “oligodT” attached magnetic beads, RNA from each sample was subjected to two rounds of enrichment for poly-A mRNAs. Upon chemical fragmentation, mRNA was converted to single-stranded cDNA. Using adapter indexes, cDNA from each library was separately barcoded. Barcoded libraries were then pooled and subsequently sequenced on a HiSeq 2000 platform (Illumina Inc., USA) using 100bp single-end sequencing runs. Sequencing was performed at the Genome Sequencing Facility of Kansas University Medical Center.

**ii) Sequence processing, alignment to reference genome, and differential gene expression analysis**

The adapter trimmer “Cutadapt” (Martin, 2011) was used to trim the adapters from sequence reads and subsequent quality filtering. Alignment of reads to *sorghum bicolor* reference genome (Sbicolor_v1.4) (Paterson et al., 2009) was conducted using the Genomic Short-read Nucleotide Alignment Program (GSNAP) (Wu and Watanabe, 2005). Differential gene expression analysis was performed using ‘DESeq2’. DESeq2 analysis was performed to test the null hypothesis of no significant two way interaction between sorghum line (2 levels: resistant and susceptible sorghum lines) and treatment (2 levels: infected with *M. phaseolina* and mock-inoculated control) for each gene within each post inoculation stage (i.e, 2, 7, and 30 DPI). A q-value (Benjamini and Hochberg, 1995) was determined for each gene to account for multiple tests. To control false discovery rate (FDR) at 5%, only the genes with q-values smaller than 0.05 were considered to be significantly differentially expressed. SorghumCyc (http://pathway.gramene.org/gramene/sorghumcyc.shtml) and Phytozome (Goodstein et al., 2012) data bases were used to identify the annotated functions of differentially expressed genes.

### B) Functional assays

**i) Plant materials, establishment, maintenance, inoculum preparation, inoculation, stalk tissue collection, preparation of tissue lysates for functional assays, and absorbance measurement**

Two charcoal rot resistant (SC599, SC35) and susceptible (Tx7000, BTx3042) sorghum genotypes were used. Plant establishment, maintenance, inoculum preparation, and inoculation were performed as described under RNASeq experiment. At 4, 7, and 10 DPI, 15 cm long stalk pieces encompassing the inoculation point were cut from five biologically replicated plants, immediately suspended in liquid nitrogen, and subsequently stored at −80 °C until used. Stalk tissues were retrieved from −80°C storage and approximately 1 g of stalk tissues (1 cm away from the symptomatic region) was quickly chopped in to liquid nitrogen (in a mortar) using a sterile scalpel. The stalk pieces were ground into a powder using a pestle. Approximately 200 mg of this tissue powder was transferred into microcentrifuge tubes filled with 1 ml of potassium phosphate buffer (50 mM potassium phosphate (pH 6.8), 0.1 mM ethylenediaminetetraacetic (EDTA), 1 mM phenylmethylsulfonyl fluoride, and 2% (wt/vol) polyvinylpolypyrrolidone (PVPP); used for all glutathione related assays), 1 ml of 1X PBS with 1mM EDTA (for catalase and peroxidase assays), and 1 ml of 1X Lysis buffer (10 mM Tris, pH 7.5, 150 mM NaCl, 0.1 mM EDTA; for superoxide dismutase assay). Buffer selections were based on the instructions by assay kit manufacturers. Samples were centrifuged at 10000 g for 10 min at 4°C. Supernatants were transferred into new microcentrifuge tubes and immediately stored at −80°C until used in assays. All absorption/fluorescence measurements were performed using a 96-well plate reader (Synergy H1 Hybrid Reader, BioTek, Winooski, VT, USA) at specified wavelengths (see below). Path length correction was performed using an option available in the plate reader during the measurements. All functional experiments were repeated twice.

**ii) Quantification of total, oxidized, and reduced glutathione concentrations**

The EnzyChrom™ GSH/GSSG Assay Kit (BioAssay Systems, Hayward, CA, USA) was used to quantify the total, oxidized, and reduced glutathione concentrations of the samples. The assay is based on an enzymatic method that utilizes Ellman’s Reagent (DTNB) and glutathione reductase (GR). DTNB reacts with glutathione to form a yellow product. The rate of change in the optical density, measured at 412 nm, is directly proportional to glutathione concentration in the sample. In the current study, the total glutathione concentration (reduced + oxidized) was determined following the protocol described by the manufacturer with some modifications. Briefly, 10μL of each sample was diluted in 90 μL 1X Assay Buffer and transferred to a Nunc™ 96-Well Polypropylene MicroWell™ Plate (Thermo Scientific Nunc, Roskilde, Denmark). The standards were prepared according to the manufacturer’s instructions. A master mix of the working reagent (WR), sufficient for all samples and standards, was prepared (105 μL 1X assay buffer, 1 μL GR enzyme, 0.25 μL NADPH and 0.5 μL DTNB per reaction). Fifty μL of WR was immediately added to each standard and sample and was well mixed. The optical density (OD) was read at 412 nm at 0 min and again at 10 min. OD_0min_ was subtracted from OD_10min_ for each standard and sample. Then, the ΔOD_BLANK_ (1X assay buffer) was subtracted from ΔOD values of all standards and the ΔΔOD’s were plotted against standard concentrations. The slope was determined using linear regression fitting and the total glutathione (GSH_TOTAL_) concentrations of the samples were calculated using the following equation:

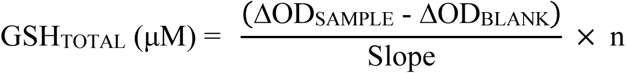

Where n = dilution factor

The same procedure explained above was used to determine the oxidized glutathione (GSSG) concentration. However, at the beginning, 45 μL from each sample was mixed with 5 μL of 1-methyl-2-vinylpyridinium triflate to scavenge GSH in the solution. From this solution, 10 μL was drawn and diluted in 90 μL 1X assay buffer to proceed further as described above. The GSSG concentrations of the samples were calculated using the following equation:

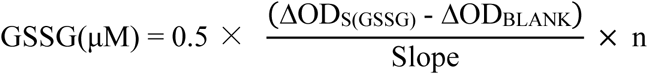

Where, ΔOD_S(GSSG)_ = sample treated with scavenger, and n = dilution factor

The reduced glutathione (GSH) concentrations of the samples were determined using the following equation:

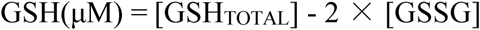

**iii) Quantification of glutathione S-transferase (GST) activity**

Glutathione S-Transferase (GST) Assay Kit (SIGMA, Saint Louis, MO, USA) was used to quantify the GST activity of samples. This assay kit utilizes 1-chloro-2,4-dinitrobenzene (CDNB) as the GSH conjugant. Upon GST-mediated conjugation of CDNB with the thiol group of GSH, there is an increase in the absorbance at 340 nm. Therefore, absorbance is directly proportional to GST-specific activity. In the current study, following the manufacturer’s instructions, a master mix containing Dulbecco’s phosphate buffered saline (19.6 mL), 200 mM L-glutathione reduced (0.2 mL), and 100 mM CDNB (0.2 mL) (sufficient for all samples) was prepared. Twenty μL of each sample was transferred to a Nunc™ 96-Well Polypropylene MicroWell™ Plate (Thermo Scientific Nunc, Roskilde, Denmark) and mixed with 180 μL of master mix. Two-hundred μL of the master mix was used as the blank. Optical density (OD) was read at 340 nm at 0 min and again at 10 min. The change in optical density (ΔOD_340_)/minute was calculated in the linear range of the plot for each sample and for the blank using the following equation:

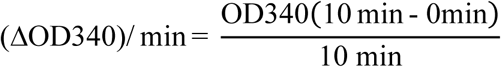

The (ΔOD_340_)/minute of the blank was subtracted from the (ΔOD_340_)/minute of the sample. This rate was used to calculate the GST-specific activity using the following equation:

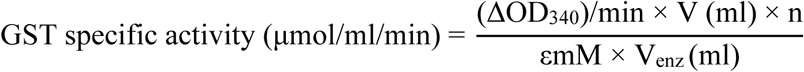

Where, n = dilution factor, εmM = extinction coefficient for CDNB conjugate at 340 nm (5.3 mM^-1^ cm^-1^), V = reaction volume (200 μL), V_enz_ = the volume of the enzyme sample tested (20 μL).

**iv) Quantification of glutathione peroxidase (GPx) activity**

The EnzyChrom™ Glutathione Peroxidase Assay Kit (EGPX-100) (BioAssay Systems, Hayward, CA, USA) was used to quantify the glutathione peroxidase activity of the samples. This assay directly measures NADPH consumption in the enzyme coupled reactions. The reduction in optical density at 340 nm is directly proportional to the enzyme activity in the sample. In the current study, following the manufacturer’s instructions, 10 μL of each standards or samples was transferred into wells of a Nunc™ 96-Well Polypropylene MicroWell™ Plate (Thermo Scientific Nunc, Roskilde, Denmark). 190 μL assay buffer was added to all standard wells. 90 μL working reagent (containing 90 μL assay buffer, 5 μL glutathione, 3 μL 35 mM NADPH and 2 μL GR enzyme per well) was quickly added to the sample/control wells and mixed briefly yet thoroughly. 100 μL of 1× substrate solution was added to all sample and control wells. Tap contents were thoroughly mixed and the optical density was immediately read at 340 nm at 0 min (OD0) and again at 4 min (OD4). OD values at 4 min were used for NADPH standards. The blank value was subtracted from the standard values and resulting ΔODs were plotted against standard concentrations to determine the slope of the standard curve. The ΔOD_S_ = (OD0 – OD4) for the samples and ΔOD_B_ = (OD0 – OD4) for the background control were determined. Finally, the GPx activity of each sample was computed using the following equation. A unit is defined as the amount of GPx that produces 1 mmole of GS-SG per min at pH 7.6 and room temperature.

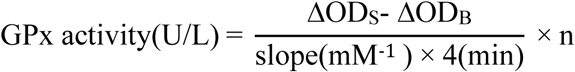

Where, n is the sample dilution factor.

**v) Quantification of glutathione reductase (GR) activity**

The EnzyChrom™ Glutathione Reductase Kit (ECGR-100) (BioAssay Systems, Hayward, CA, USA) was used to quantify the glutathione reductase activity of the samples. This assay utilizes Ellman’s method in which DTNB reacts with the GSH generated from the reduction of GSSG by the GR in a sample to form a yellow product (TNB^2-^). The rate of change in optical density, measured at 412 nm, is directly proportional to GR activity in the sample. In the current study, following manufacturer’s instruction, 20 μL from each sample, 100 μL of calibrator and 100 μL assay buffer were transferred to separate wells in a Nunc™ 96-well polypropylene MicroWell™ plate (Thermo Scientific Nunc, Roskilde, Denmark). Eighty μL of working reagent (containing 8 μL substrate, 8 μL co-substrate, 1 μL GDH, 0.5 μL DTNB and 70 μL assay buffer per well) was added to each sample well and mixed. The plate was incubated at 25°C for 10 min and the optical density was read at 412 nm at 10 min and again at 30 min. OD10 was subtracted from OD30 for each sample to compute the ΔOD_S_. GR activity was calculated using the equation below. A Unit (U) of GR is the defined as the amount of GR that will catalyze the conversion of 1 μmole of GSSG to 2 μmole GSH per min at pH 7.6.

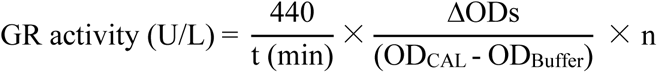

Where, OD_CAL_ and OD_Buffer_ are OD412 nm (OD0) values of the calibrator and assay buffer; and t is the reaction time (20 min), and n is the dilution factor.

### C) Assessment of the impact of exogenous glutathione application on charcoal rot disease severity

**i) Establishment and maintenance of plants**

A greenhouse experiment was conducted with two charcoal-rot-resistant (SC599, SC35) and two susceptible (Tx7000, BTx3042) sorghum lines. The experiment was arranged in randomized complete block design (RCBD) with three blocks. The seeds treated with captan (N-trychloromethyl thio-4-cyclohexane-1,2 dicarboxamide) were planted in 19 L Poly Tainer pots filled with Metro-Mix 360 growing medium (Sun Gro Bellevue, WA, U.S.A) and kept in a greenhouse at 25-32°C with a 16-h light/8-h dark photoperiod. Two weeks after seedling emergence, each pot was thinned to three seedlings. There were three pots per genotype per block and pots were randomly assigned for three inoculation treatments (pathogen, pathogen + glutathione, and mock-inoculated control), respectively. The treatment structure was a 4 × 3 factorial where factors consisted of four sorghum genotypes and three inoculation treatments. The three plants in each pot were considered as sub sample units and their averages were used for final data analysis. The experiment was repeated twice.

**ii) Inoculum preparation, inoculation, glutathione application, and measurement of disease severity**

Inoculum preparation and inoculation were performed as described under RNASeq experiment. A 10 mM L-GSH (reduced glutathione) (SIGMA, Saint Louis, MO, USA) solution was prepared by dissolving glutathione in sterile distilled water. At 5 and 10 days post inoculation (DPI), 0.1 mL of glutathione solution was injected into 3 plants in each pot assigned for pathogen + glutathione treatment using the same point as for inoculations. Plants in pots that were assigned for pathogen and mock inoculation treatments were injected with 0.1 mL of sterile distilled water at same days (5 and 10 DPI). All plants were harvested at 35 d after initial inoculation. Stems were split longitudinally to measure the disease severity, lesion length (cm).

### D) Statistical analysis of functional and disease severity data

Data were analyzed for variance (ANOVA) using the PROC GLIMMIX procedure of SAS software version 9.2 (SAS Institute, 2008). Variance components for fixed factors were estimated using restricted maximum likelihood (REML) method. The genotype and inoculation treatment were considered fixed while repeated experiments and block were treated as random. Studentized residual plots and Q-Q plots were used to test the assumptions of identical and independent distribution of residuals and their normality, respectively. Whenever heteroskedasticity was observed, appropriate heterogeneous variance models were fitted to meet the model assumptions by specifying a random/group statement (group = genotype or inoculation treatment) following the model statement. Bayesian information criterion (BIC) was used to determine the most parsimonious model. Means separations were carried out using the PROC GLMMIX procedure of SAS. Main effects of factors were determined with adjustments for multiple comparisons using the Tukey-Kramer test. Whenever the genotype × treatment interaction was statistically significant, the simple effects of inoculation treatment were determined at each genotype level.

## ACKNOWLEDGEMENTS

The Kansas Grain Sorghum Commission is gratefully acknowledged for their financial support of this research. This paper is Contribution No. 20-###-J from the Kansas Agricultural Experiment Station, Manhattan.

---

**Supplementary table 1.** Glutathione related genes that are differentially expressed between charcoal rot resistant (SC599) and susceptible (Tx7000) sorghum genotypes in response to Macrophomina phaseolina inoculation at 2^nd^ and 7^th^ days post inoculation.

| Gene name | Gene annotation | Geno × Trt*<br>q-value | S599 (MP-CON)† |  | Tx7000 (MP-CON) |  |
| --- | --- | --- | --- | --- | --- | --- |
|  |  |  | log2 DE‡ | q-value | log2 DE | q-value |
| 2 days post inoculation |  |  |  |  |  |  |
| *Sb01g001130* | Glutathione S-transferase | 1.1E-02 | 0.03 | 9.7E-01 | -1.90 | 4.9E-03 |
| *Sb08g006680* |  | 2.4E-02 | 0.04 | 9.5E-01 | -1.13 | 2.9E-02 |
| *Sb01g030800* |  | 7.0E-04 | 0.99 | 2.0E-02 | -0.90 | 4.1E-02 |
| *Sb03g044980* |  | 1.4E-02 | 1.61 | 1.9E-06 | 0.35 | 2.7E-01 |
| *Sb01g031000* |  | 5.3E-03 | -1.17 | 1.6E-01 | 1.51 | 6.5E-02 |
| *Sb02g027080* |  | 3.3E-02 | 0.51 | 2.4E-01 | 1.88 | 9.8E-02 |
| *Sb03g045840* |  | 4.3E-02 | 0.07 | 9.6E-01 | 2.18 | 2.2E-04 |
| *Sb03g045830* |  | 1.2E-03 | -0.28 | 7.8E-01 | 2.18 | 3.2E-05 |
| *Sb04g032520* | Glutathione peroxidase | 3.9E-07 | 0.99 | 8.1E-05 | -0.46 | 1.5E-01 |
| *Sb03g000550* | Glutaredoxin | 1.8E-04 | -1.47 | 3.0E-07 | 0.43 | 3.6E-01 |
| *Sb02g041880* |  | 6.4E-05 | -0.71 | 9.0E-03 | 0.84 | 2.5E-02 |
| 7 days post inoculation |  |  |  |  |  |  |
| *Sb03g025210* | Glutathione S-transferase | 3.7E-05 | 0.63 | 8.3E-01 | -5.18 | 4.7E-11 |
| *Sb01g030800* |  | 3.0E-11 | 1.55 | 2.1E-02 | -2.41 | 6.4E-11 |
| *Sb01g030810* |  | 6.9E-03 | 0.91 | 6.2E-01 | -2.06 | 6.0E-05 |
| *Sb08g007310* |  | 3.5E-02 | -0.85 | 9.6E-01 | -1.32 | 3.0E-03 |
| *Sb06g017640* |  | 1.8E-02 | -0.17 | 9.3E-01 | -1.17 | 1.4E-04 |
| *Sb09g001690* |  | 8.5E-03 | 1.00 | 3.1E-01 | -0.94 | 2.5E-01 |
| *Sb10g008310* |  | 1.9E-04 | 0.49 | 4.4E-01 | -0.69 | 1.8E-02 |
| *Sb09g003700* |  | 4.2E-02 | -1.42 | 1.7E-01 | 0.29 | 6.8E-01 |
| *Sb06g017110* |  | 8.9E-03 | -0.50 | 4.9E-01 | 0.52 | 1.4E-01 |
| *Sb09g003750* |  | 5.0E-04 | - | - | 0.69 | 2.1E-04 |
| *Sb03g015070* |  | 4.3E-02 | - | - | 0.81 | 6.7E-01 |
| *Sb04g023210* |  | 2.8E-02 | -0.25 | 9.0E-01 | 0.93 | 1.5E-03 |
| *Sb05g007005* |  | 2.4E-04 | -0.64 | 3.3E-01 | 1.03 | 2.2E-03 |
| *Sb09g003690* |  | 1.2E-06 | -1.22 | 4.1E-02 | 1.05 | 4.2E-05 |
| *Sb01g005990* |  | 3.1E-02 | -0.25 | 9.4E-01 | 1.40 | 1.4E-06 |
| *Sb01g001130* |  | 2.0E-02 | -0.35 | 8.8E-01 | 1.46 | 3.2E-03 |
| *Sb01g006010* |  | 7.5E-03 | -0.16 | 9.7E-01 | 2.07 | 8.4E-07 |
| *Sb03g045840* |  | 8.9E-04 | -1.24 | 4.3E-01 | 2.16 | 5.3E-05 |
| *Sb08g006690* |  | 1.2E-05 | -1.07 | 2.7E-01 | 2.17 | 5.7E-07 |
| *Sb05g001525* |  | 3.6E-02 | -0.90 | 7.3E-01 | 2.24 | 8.0E-03 |
| *Sb01g031030* |  | 7.8E-05 | -1.03 | 4.2E-01 | 2.44 | 7.4E-09 |
| *Sb08g007300* |  | 1.1E-04 | - | 4.2E-01 | 2.61 | 4.2E-07 |
| *Sb01g030930* |  | 4.8E-05 | -0.04 | - | 3.23 | 3.6E-16 |
| *Sb01g006000* |  | 6.2E-08 | -0.11 | 9.8E-01 | 3.35 | 5.3E-17 |
| *Sb02g027080* |  | 3.7E-09 | -0.32 | 8.6E-01 | 3.79 | 1.6E-09 |
| *Sb01g030870* |  | 6.2E-04 | -0.06 | 9.9E-01 | 3.83 | 3.0E-10 |
| *Sb01g030790* |  | 1.3E-02 | 0.09 | 9.9E-01 | 3.90 | 2.8E-07 |
| *Sb03g031780* |  | 6.5E-04 | -0.28 | 9.5E-01 | 4.04 | 2.9E-11 |
| *Sb01g030880* |  | 5.2E-05 | -1.51 | 3.6E-01 | 4.14 | 9.4E-12 |
| *Sb03g045830* |  | 4.3E-09 | -0.97 | 4.4E-01 | 4.18 | 1.3E-13 |
| *Sb04g022250* |  | 1.6E-04 | -0.01 | 1.0E+00 | 4.59 | 6.0E-26 |
| *Sb01g030980* |  | 2.1E-20 | -0.04 | 9.9E-01 | 5.58 | 7.1E-89 |
| *Sb01g030830* |  | 5.5E-03 | 0.70 | 8.4E-01 | 5.99 | 6.7E-06 |
| *Sb01g031040* |  | 2.4E-07 | - | - | 6.58 | 2.3E-15 |
| *Sb01g030990* |  | 5.2E-17 | -0.04 | - | 6.83 | 4.5E-08 |
| *Sb01g031020* |  | 3.1E-03 | - | - | 7.17 | 9.4E-09 |
| *Sb01g031000* |  | 2.5E-35 | -2.42 | 1.1E-03 | 7.24 | 2.8E-40 |
| *Sb01g031010* |  | 5.8E-14 | 0.07 | - | 7.48 | 3.9E-10 |
| *Sb02g003090* |  | 2.3E-20 | -0.94 | 6.2E-01 | 7.88 | 2.1E-89 |
| *Sb01g031050* |  | 4.0E-06 | - | - | 8.00 | 1.3E-11 |
| *Sb02g038130* |  | 2.3E-34 | -0.20 | - | 9.54 | 1.0E-35 |
| *Sb08g016750* | Glutathione synthetase | 2.4E-06 | -0.92 | 6.2E-02 | 1.92 | 1.2E-05 |
| *Sb09g002470* | Glutamate cysteine ligase | 1.5E-03 | -0.88 | 2.5E-01 | 0.85 | 6.5E-03 |
| *Sb10g005820* | Glutathione peroxidase | 1.6E-04 | 0.83 | 3.8E-01 | -2.04 | 1.8E-05 |
| *Sb06g024920* |  | 8.5E-04 | -0.24 | 8.8E-01 | 0.99 | 1.3E-07 |
| *Sb01g034870* |  | 5.9E-07 | -0.83 | 1.0E-01 | 1.28 | 1.5E-05 |
| *Sb04g032520* |  | 5.1E-03 | 0.19 | 9.6E-01 | 2.13 | 1.1E-10 |
| *Sb01g035940* |  | 1.6E-03 | - | - | 6.64 | 1.5E-07 |
| *Sb04g036870* | Glutathione reductase | 1.5E-03 | -0.26 | 9.0E-01 | 1.52 | 1.4E-07 |
| *Sb01g021980* |  | 6.1E-05 | -0.33 | 9.2E-01 | 2.95 | 7.6E-10 |
| *Sb03g000550* | Glutaredoxin | 1.8E-05 | 1.28 | 3.4E-01 | -3.54 | 4.8E-11 |
| *Sb06g014830* |  | 1.2E-02 | 0.28 | 8.8E-01 | -1.03 | 8.4E-03 |

\* Geno × Trt = genotype by treatment interaction where treatment consists of *M. phaseolina* and Control inoculations. †MP = *M. phaseolina*, CON = control. ‡ log2 DE = log2 fold differential expression.

## REFERENCES

1. Agrawal, G.K., Rakwal, R., Jwa, N.S. and Agrawal, V.P. (2002) Effects of signaling molecules, protein phosphatase inhibitors and blast pathogen (*Magnaporthe grisea*) on the mRNA level of a rice (*Oryza sativa* L.) phospholipid hydroperoxide glutathione peroxidase (OsPHGPX) gene in seedling leaves. Gene, 283, 227–236.

2. Airaki, M., Sánchez-Moreno, L., Leterrier, M., Barroso, J. B., Palma, J. M. and Corpas, F. J. (2011) Detection and quantification of S-nitrosoglutathione (GSNO) in pepper (*Capsicum annuum* L.) plant organs by LC-ES/MS. Plant Cell Physiol. 52, 2006–2015.

3. Alscher, R.G and Hess, J.L. (1993) Antioxidants in Higher Plants. CRC press, Boca Raton, FL.

4. Bandara, Y.M.A.Y., Tesso, T.T., Bean, S.R., Dowell, F.E. and Little, C.R. 2017 (a). Impacts of fungal stalk rot pathogens on physicochemical properties of sorghum grain. Plant Dis. https://doi.org/10.1094/PDIS-02-17-0238-RE.

5. Bandara, Y.M.A.Y., Weerasooriya, D.K., Tesso, T.T., Prasad, P.V.V. and Little, C.R. 2017 (b). Stalk rot fungi affect grain sorghum yield components in an inoculation stage-specific manner. Crop Prot. 94, 97–105.

6. Bandara, Y.M.A.Y., Weerasooriya, D.K., Tesso, T.T. and Little, C.R. 2017 (c). Stalk rot diseases impact sweet sorghum biofuel traits. BioEnergy Res. 10, 26–35.

7. Bandara, Y.M.A.Y., Weerasooriya, D.K., Tesso, T.T. and Little, C.R. 2016. Stalk rot fungi affect leaf greenness (SPAD) of grain sorghum in a genotype- and growth stage-specific manner. Plant Dis. 100, 2062–2068.

8. Bandara, YMAY., Perumal, R. and Little C. R. (2015) Integrating resistance and tolerance for improved evaluation of sorghum lines against Fusarium stalk rot and charcoal rot. Phytoparasitica 43, 485–499.

9. Bartling, D., Radzio, R., Steiner, U. and Weiler E.W. (1993) A glutathione S-transferase with glutathione peroxidase activity from *Arabidopsis thaliana*: molecular cloning and functional characterization. Eur. J. Biochem. 216, 579–586.

10. Benjamini, Y. and Hochberg, Y. (1995) Controlling the false discovery rate: a practical and powerful approach to multiple testing. Journal of the Royal Statistical Society: Series B (Methodological) 57, 289–300.

11. Berhane, K., Widersten, M., Engstrom, A., Kozarich, J. and Mannervik, B. (1994) Detoxication of base propenals and other alpha, beta-unsaturated aldehyde products of radical reactions and lipid peroxidation by human glutathione S-transferases. Proc. Natl. Acad. Sci. USA, 91, 1480–1484.

12. Bianchi, M.W., Roux, C. and Vartanian, N. (2002) Drought regulation of GST8, encoding the Arabidopsis homologue of ParC/Nt107 glutathione transferase/peroxidase, Physiol. Plant. 116, 96–105.

13. Coleman, J.O.D., Blake-Kalff, M.M.A. and Davies T.G.E. (1997) Detoxification of xenobiotics by plants: chemical modification and vacuolar compartmentation. Trends Plant Sci. 2, 144–151.

14. Creissen, G., Firmin, J., Fryer, M., Kular, B., Leyland, N., Reynolds, H., Pastori, G., Wellburn, F., Baker, N., Wellburn, A. and Mullineaux, P. (1999) Elevated glutathione biosynthetic capacity in the chloroplasts of transgenic tobacco plants paradoxically causes increased oxidative stress. Plant Cell, 11, 1277–1291.

15. Cummins, I., Cole, D.J. and Edwards, R. (1999) A role for glutathione transferases functioning as glutathione peroxidases in resistance to multiple herbicides in black-grass. Plant Journal, 18, 285–292.

16. Danielson, U.H., Esterbauer, H. and Mannervik, B. (1987) Structure-activity relationships of 4-hydroxyalkenals in the conjugation catalyzed by mammalian glutathione S-transferases. Biochem. J. 247, 707–712.

17. Dixon, D.P., Cummins, I., Cole, D.J. and Edwards, R. (1998) Glutathione-mediated detoxification systems in plants. Curr Opin Plant Biol. 1, 258–266.

18. Droge, W. (2002) Free radicals in the physiological control of cell function. Physiol Rev. 82, 47–95.

19. Dron, M., Clouse, S.D., Dixon, R.A., Lawton, M.A. and Lamb, C.J. (1988) Glutathione and fungal elicitor regulation of a plant defense gene promoter in electroporated protoplasts. Proc Natl Acad Sci USA, 85, 6738–6742.

20. Edwards, R., Blount, J.W. and Dixon, R.A. (1991) Glutathione and elicitation of the phytoalexin response in legume cell cultures. Planta, 184, 403–409.

21. Edwards, R., Dixon, D.P. and Walbot, V. (2000) Plant glutathione S-transferases: enzymes with multiple functions in sickness and in health. Trends Plant Sci. 5, 193–198.

22. El-Zahaby, H.M., Gullner, G. and Kiraly, Z. (1995) Effects of powdery mildew infection of barley on the ascorbate–glutathione cycle and other antioxidant in different host–pathogen interactions. Phytopathology, 85, 1225–1230.

23. Fernandez, J. and Wilson, R.A. 2014. Characterizing roles for the glutathione reductase, thioredoxin reductase and thioredoxin peroxidase-encoding genes of *Magnaporthe oryzae* during rice blast disease. PLoS One 9: e87300.

24. Foyer, C.H., Lelandais, M. and Kunert, K.J. (1994) Photooxidative stress in plants. Physiol. Plant. 92, 696–717.

25. Foyer, C.H. and Noctor, G. (2003) Redox sensing and signalling associated with reactive oxygen in chloroplasts, peroxisomes and mitochondria. Physiol. Plant. 119, 355– 364.

26. Foyer, C.H. and Noctor, G. (2005) Redox homeostasis and antioxidant signaling: a metabolic interface between stress perception and physiological responses. Plant Cell, 17, 1866–1875.

27. Foyer, C.H. and Noctor, G. (2009) Redox regulation in photosynthetic organisms: signaling, acclimation, and practical implications. Antioxid. Redox Signal. 11, 861–905.

28. Foyer, C.H. and Noctor, G. (2011) Ascorbate and glutathione: the heart of the redox hub. Plant Physiol. 155, 2–18.

29. Gomez, L.D., Noctor, G., Knight, M. and Foyer, C.H. (2004) Regulation of calcium signaling and gene expression by glutathione. J Exp Bot. 55, 1851–1859.

30. Gonnen, M.V. and Schlösser, E. (1993) Oxidative stress in interaction between *Avena sativa* L. and *Drechlera* spp.. Physiol Mol Plant Path. 42, 221–234.

31. Goodstein, D.M., Shu, S., Howson, R., Neupane, R., Hayes, R.D., Fazo J. and Rokhsar D.S. (2012) Phytozome: a comparative platform for green plant genomics. Nucleic Acids Research 40, 1178–1186.

32. Hatier, J. H. B. and Gould, K. S. (2008) Foliar anthocyanins as modulators of stress signals. J. Theor. Biol. 253, 625–627.

33. Huber, P. C., Almeida, W. P. and Fátima, Â. D. (2008) Glutathione and related enzymes: biological roles and importance in pathological processes. Química Nova, 31, 1170–1179.

34. Islam, M.S., Haque, M.S., Islam, M.M., Emdad, E.M., Halim, A., Hossen, Q.M.M., Hossain, M.Z., Ahmed, B., Rahim, S., Rahman, M.S. and Alam, M.M. 2012. Tools to kill: genome of one of the most destructive plant pathogenic fungi *Macrophomina phaseolina*. BMC Genomics 13: 1.

35. Kiyosue, T., Yamaguchi-Shinozaki, K. and Shinozaki, K. (1993) Characterization of two cDNAs (ERD11 and ERD13) for dehydration-inducible genes that encode putative glutathione S-transferases in *Arabidopsis thaliana*. FEBS Lett. 335, 189–192.

36. Kuzniak, E. and Sklodowska, M. (1999) The effect of Botrytis cinerea infection on ascorbate glutathione cycle in tomato leaves. Plant Science, 148, 69–76.

37. Levine, A., Tenhaken, R., Dixon, R. and Lamb, C. (1994) H_2_O_2_ from the oxidative burst orchestrates the plant hypersensitive disease resistance response. Cell, 79, 583–593.

38. Liao, W., Ji, L., Wang, J., Chen, Z., Ye, M., Ma, H. and An, X. (2014) Identification of glutathione S-transferase genes responding to pathogen infestation in *Populus tomentosa*. Funct Integr Genomics, 14, 517–529.

39. Margis, R., Dunand, C., Teixeira, F. K. and Margis-Pinheiro, M. (2008) Glutathione peroxidase family–an evolutionary overview. FEBS journal, 275, 3959–3970.

40. Marí, M., Morales, A., Colell, A., García-Ruiz, C. and Fernández-Checa, J. C. (2009) Mitochondrial glutathione, a key survival antioxidant. Antioxid Redox Signal. 11, 2685–2700.

41. Marrs, K.A., Alfenito, M.R., Lloyd, A.M. and Walbot, V. (1995) A glutathione S-transferase involved in vacuolar transfer encoded by the maize gene Bronze-2. Nature, 375, 397–400.

42. Martin, M. (2011) Cutadapt removes adapter sequences from high-throughput sequencing reads. EMBnet Journal 17, 10–12.

43. Martinoia, E., Grill, E., Tommasini, R., Kreuz, K. and Amrhein, N. (1993) ATP-dependent glutathione S-conjugate ‘export’ pump in the vacuolar membrane of plants. Nature, 364, 247–249.

44. Matern, U., Reichenbach, C. and Heller, W. (1986) Efficient uptake of flavonoids into parsley (*Petroselinum hortense*) vacuoles requires acylated glycosides. Planta, 167, 183–189.

45. Mauch, F. and Dudler, R. (1993) Differential induction of distinct glutathione-S-transferases of wheat by xenobiotics and by pathogen attack. Plant Physiol. 102, 1193–201.

46. Meloni, D. A., Oliva, M. A., Martinez, C. A. and Cambraia, J. (2003) Photosynthesis and activity of superoxide dismutase, peroxidase and glutathione reductase in cotton under salt stress. Environ Exper Bot. 49, 69–76.

47. Mittler, R., Vanderauwera, S., Gollery, M. and Van Breusegem, F. (2004). Reactive oxygen gene network of plants. Trends Plant Sci. 9, 490–498.

48. Moons, A. (2003) Osgstu3 and osgtu4, encoding tau class glutathione S-transferases, are heavy metal- and hypoxic stress-induced and differentially salt stress responsive in rice roots, FEBS Lett. 553, 427–432.

49. Nimse, S.B. and Pal, D. (2015) Free radicals, natural antioxidants, and their reaction mechanisms. RSC Advances. 5, 7986–8006.

50. Noctor, G. and Foyer, C.H. (1998) Ascorbate and glutathione: keeping active oxygen under control. Annu. Rev. Plant Physiol. Plant Mol. Biol. 49, 249–279.

51. Paterson, A.H., Bowers, J.E., Bruggmann, R., Dubchak, I., Grimwood, J., Gundlach, H. and Poliakov, A. (2009). The *Sorghum bicolor* genome and the diversification of grasses. Nature 457, 551–556.

52. Sandermann, H. (1992) Plant metabolism of xenobiotics. Trends Biol. Sci. 17, 82–84.

53. Sandermann, H. (1994) Higher plant metabolism of xenobiotics: the “green liver” concept. Pharmacogenetics, 4, 225–241.

54. Seppanen, M.M., Cardi, T., Hyokki, M.B. and Pehu, E. (2000) Characterisation and expression of cold induced glutathione S-transferase in freezing tolerant *Solanum commersonii*, sensitive *S. tuberosum* and their interspecific somatic hybrids, Plant Sci. 153, 125–133.

55. Shahidi, F. and Zhong, Y. (2010). Novel antioxidants in food quality preservation and health promotion. Eur J Lipid Sci Technol. 112, 930–940.

56. Tarr, SAJ. 1962. Root and stalk diseases: Red stalk rot, Colletotrichum rot, anthracnose, and red leaf spot, in: Diseases of Sorghum, Sudan Grass and Brown Corn. Commonwealth Mycological Institute, Kew, Surrey, UK, pp. 58–73.

57. Tesso, T., Claflin, L.E. and Tuinstra MR. (2004) Estimation of combining ability for resistance to Fusarium stalk rot in grain sorghum. Crop Sci. 44, 1195–1199.

58. Tesso, T., Perumal, R., Little, C.R., Adeyanju, A., Radwan, G.L., Prom, L.K. and Magill, C.W. (2012) Sorghum pathology and biotechnology-a fungal disease perspective: Part II. Anthracnose, stalk rot, and downy mildew. Eur J Plant Pathol. 6, 31–44.

59. Vanacker, H., Carver, T.L.W. and Foyer, C.H. (1998) Pathogen-induced changes in the antioxidant status of the apoplast in barley leaves, Plant Physiol. 117, 1103–1114.

60. Wolf, A.E., Dietz, K.J. and Schroder, P. (1996) A carboxypeptidase degrades glutathione conjugates in the vacuoles of higher plants. FEBS Lett. 384, 31–34.

61. Xiang, C. and Oliver, D. J. (1998) Glutathione metabolic genes coordinately respond to heavy metals and jasmonic acid in Arabidopsis. Plant Cell, 10, 1539–1550.

